# Optical Conductivity as a Stage-Sensitive Imaging Biomarker for Alzheimer’s Disease

**DOI:** 10.64898/2026.06.02.729558

**Authors:** Hao Yang, Shixie Jiang, Huabei Jiang

## Abstract

Early detection of Alzheimer’s disease (AD) remains limited by the inability of current imaging biomarkers to capture dynamic tissue changes preceding overt neurodegeneration. Conventional approaches primarily reflect molecular burden or structural atrophy and incompletely characterize the intermediate microenvironmental remodeling linking early pathology to clinical progression. Here, we introduce optical conductivity as a novel imaging biomarker characterizing tissue responses to optical-frequency electromagnetic fields. Using an optical wave tomography (OWT) framework, optical conductivity is reconstructed from diffuse optical measurements by integrating absorption-driven energy deposition with microstructure-dependent field redistribution. Applied to an APP/PS1 mouse model across young and old cohorts, optical conductivity revealed a significant age-by-disease interaction (F = 9.20, p = 0.0071). Quantitatively, its value was reduced in young AD animals relative to controls (∼13%; Cohen’s d = 1.93) but showed attenuation or reversal in older animals, indicating a stage-dependent crossover. Consistently, optical conductivity achieved strong classification performance in the young cohort (AUC = 0.80), outperforming absorption and exceeding scattering, with improved performance via multimodal integration (AUC = 0.87). Together, these findings suggest that optical conductivity captures stage-dependent microenvironmental remodeling not detectable by conventional optical parameters, offering potential for early detection and disease staging.

## 2. Introduction

Alzheimer’s disease (AD) is the leading cause of dementia and a major global health burden, affecting over 55 million individuals worldwide [1, 2]. A defining feature of AD is its prolonged preclinical and early-stage phase, during which pathological processes such as amyloid-β accumulation, tau pathology, synaptic dysfunction, and microstructural tissue remodeling evolve years before the onset of clinical symptoms [3-5]. This extended window represents a critical opportunity for early detection and intervention; however, current diagnostic approaches remain limited in their ability to capture these early and dynamic changes [6-9].

In clinical practice, AD diagnosis has advanced to follow a biomarker-driven framework integrating cognitive assessment with molecular and imaging markers. Structural magnetic resonance imaging (MRI) is used to assess neurodegeneration [10, 11], while amyloid and tau positron emission tomography (PET) provide direct evidence of pathological depositions [12-14]. Cerebrospinal fluid (CSF) and emerging plasma biomarkers, including phosphorylated tau species, offer minimally invasive measures of underlying pathology [15-18]. Together, these modalities define the A/T/N classification system linking clinical presentation with amyloid (A), tau (T), and neurodegeneration (N) [19-21].

Despite these advances, the current framework remains limited in its ability to resolve early-stage disease and predict progression. Biomarker evaluation is often initiated after cognitive symptoms emerge, yet early symptoms are frequently nonspecific and may not correspond to underlying pathology. In asymptomatic individuals, biomarker positivity does not necessarily indicate imminent decline, and many individuals remain clinically stable for extended periods. More fundamentally, existing biomarkers primarily reflect molecular accumulation or macroscopic structural loss, rather than the intermediate microenvironmental changes that link early pathology to functional impairment. As a result, a critical gap remains in the ability to detect and characterize stage-dependent tissue remodeling during the transition from preclinical to symptomatic disease.

Neuroimaging approaches provide spatially resolved information that complements molecular biomarkers, but current modalities remain limited in early-stage disease. MRI primarily reflects macroscopic atrophy that emerges after substantial neuronal loss, while PET imaging detects amyloid and tau depositions without resolving their downstream structural or functional consequences. These limitations highlight the need for imaging approaches capable of capturing early, dynamic changes in tissue organization. Optical imaging techniques, including near-infrared spectroscopy (NIRS) [22-25] and diffuse optical tomography (DOT) [26-29], offer a noninvasive and scalable approach to probing tissue properties in vivo. Particle-like optical parameters measured by NIRS and DOT, including the absorption coefficient (*μ*_*a*_) and reduced scattering coefficient 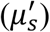, reflect chromophore concentration and microstructural heterogeneity, respectively. However, these parameters represent partially decoupled physiological dimensions and do not provide a unified description of how electromagnetic wave is distributed and dissipated within tissue.

Emerging evidence suggests that early-stage AD involves subtle microenvironmental alterations, including changes in cellular organization, membrane structure, extracellular composition, and local water distribution [30-34]. These changes are expected to influence the interaction of electromagnetic wave with tissue. An imaging metric that captures this integrated wave-tissue interaction may therefore provide improved sensitivity to early and stage-dependent tissue reorganization.

In this work, we introduce optical conductivity as a novel imaging biomarker that characterizes tissue response to optical-frequency electromagnetic waves through a power-loss–based formulation. Using an optical wave tomography (OWT) framework, optical conductivity as a wave nature-associated parameter is reconstructed from diffuse optical measurements by integrating absorption-driven energy deposition with microstructure-dependent field redistribution. Unlike NIRS and DOT, OWT provides an integrated, field-based representation of tissue organization. We hypothesize that optical conductivity is sensitive to stage-dependent microenvironmental changes that are not detectable using absorption or scattering alone. To test this hypothesis, we applied OWT to an Alzheimer’s disease mouse model across distinct age groups. We demonstrate that optical conductivity reveals a significant age-dependent disease interaction, exhibiting a stage-specific crossover pattern that distinguishes early and later disease states. These findings suggest that optical conductivity provides a complementary and biologically informative imaging metric with potential relevance for early detection and disease staging.

## 3. Methods and Materials

### 3.1 Theoretical Basis of Optical Wave Tomography

Biological tissue interacting with optical-frequency electromagnetic waves can be described from two complementary perspectives. In classical electromagnetic wave optics, tissue is treated as a lossy dielectric medium, and its dissipative response under an oscillating electric field is commonly characterized by an effective conductivity at optical frequency, which quantifies the rate of electromagnetic energy conversion into heat [35-37]. In diffuse optics, light is treated as classic particles/photons and tissue–photon interaction is then described macroscopically by the absorption coefficient *μ*_*a*_ and the reduced scattering coefficient 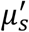 where *μ*_*a*_ represents molecular absorption (energy loss) and 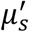 captures microstructural heterogeneity that redistributes the photon density and modulates photon residence time [38-41]. Although formulated differently, both frameworks ultimately describe optical energy attenuation and dissipation inside tissue. These two formulations therefore describe the same physical process (i.e., local energy dissipation) from a complementary perspective.

A direct conceptual bridge between the two different descriptions can be established through volumetric power-loss density. In electromagnetics, the time-averaged dissipated power per unit volume can be expressed as [35]

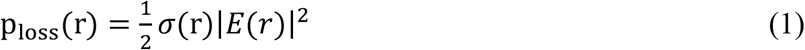

where σ denotes an effective optical-frequency conductivity and *E* is the local electric field. Under diffuse light conditions, the optical fluence rate Φ is proportional to the ensemble-averaged electromagnetic energy density, providing a macroscopic description of the underlying microscopic electric field distribution [38, 39]. In diffuse optics, the absorbed power density is [38, 40]

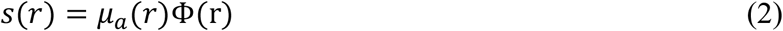

where Φ is the photon density or optical fluence rate (energy flux per unit area). Because both *p*_*loss*_ and *s* represent the same local optical energy dissipation, this correspondence provides a conceptual basis for us to obtain an effective optical conductivity metric from the DOT-derived optical parameters (i.e., *μ*_*a*_ and Φ) and computation of electric field.

Importantly, the “optical conductivity” discussed here as an optical wave nature-associated parameter should not be interpreted as low-frequency electrical conductivity associated with ionic charge transport. Rather, it is an exploratory, effective descriptor of optical-frequency energy loss [42, 43]. Since dissipation in tissue is shaped not only by absorption but also by microstructure-dependent field redistribution (reflected by 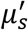 and dielectric properties), this metric is expected to capture physical features not fully characterized by *μ*_*a*_ or 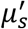 alone and even 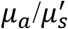combined. Biologically, changes in tissue composition and microstructure, such as alterations in cellular density, membrane organization, molecular polarizability, and structural disorder, may influence this integrated dissipative response, making optical conductivity a potentially informative composite imaging marker in vivo.

### 3.2 Reconstruction Algorithm for Optical Conductivity

The reconstruction of optical conductivity proceeds in two conceptually distinct stages: (1) estimation of conventional optical properties to determine energy deposition, and (2) recovery of conductivity through electromagnetic field modeling. In the first stage, DOT was used to recover the spatial distributions of the absorption coefficient *μ*_*a*_(*r*) and reduced scattering coefficient 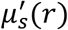. These reconstructed optical properties were subsequently employed in a forward photon diffusion calculation under planar surface illumination to compute the internal photon density Φ(r). The absorption and photon density distributions were then combined to form a power loss density term defined as *s*(*r*) = *μ*_*a*_(*r*)Φ(r), which provides the quantitative link between optical transport modeling and conductivity recovery. In the second stage, the optical conductivity σ(*r*) was estimated iteratively by separating the nonlinear product relationship *s*(*r*) = σ(*r*)|*E*(*r*)|^2^ through repeated finite-element solutions to a scalar Helmholtz equation. Starting from an initial guess of σ(*r*), the electromagnetic field distribution was computed, the predicted power loss density σ(*r*)|*E*(*r*)|^2^ was evaluated, and σ(*r*) was updated accordingly. This process was repeated until the discrepancy between the DOT-derived *s*(*r*) and the computationally modeled σ(*r*)|*E*(*r*)|^2^ reached convergence within a prescribed tolerance.

Specifically, under the diffusion approximation, photon transport in tissue is governed by the photon diffusion equation with Robin boundary conditions [29]:

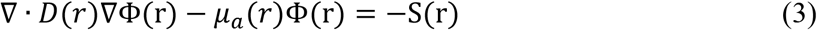

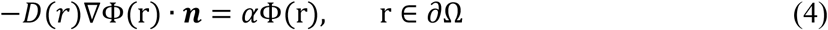

In these equations, Φ(r) denotes the photon fluence rate at position r, and S(r) represents the optical source term. The diffusion coefficient is defined as 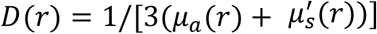. The boundary condition in Eq. (4) is a Robin boundary condition, where ***n*** denotes the outward unit normal vector on the boundary ∂Ω. The parameter *α* accounts for refractive index mismatch at the tissue boundary and models partial photon escape. The optical property distributions were updated using a regularized Newton-type method [29]:

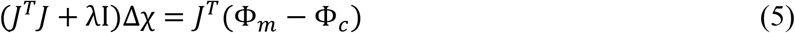

where Φ_*m*_ and Φ_*c*_ denote measured and computed boundary data, J is the Jacobian matrix of the measured boundary photon density with respect to the optical parameters (*μ*_*a*_ and 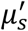), λ is the regularization parameter, and Δχ represents the update vector of the optical property profiles. After reconstruction of *μ*_*a*_(*r*) and 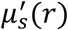, these distributions were substituted back into the diffusion model, and the forward problem was solved again under planar surface illumination to compute Φ(r). Following the energy-loss formulation described in Eq. (2), the internal optical power-loss density was evaluated as *s*(*r*) = *μ*_*a*_(*r*)Φ(r), which represents the spatial distribution of optical energy dissipation within tissue. In the reconstruction process, this loss density served as the quantitative intermediate variable linking the DOT-derived optical parameters to the subsequent conductivity estimation.

Conductivity reconstruction was performed by solving the following scalar Helmholtz equation for the electromagnetic field in finite-element form [44, 45]:

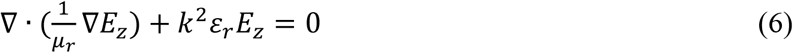

where *E*_*z*_ denotes the electric-field component used in the model, *μ*_*r*_ and *ε*_*r*_ are the relative permeability and permittivity, and k is the wave number. The Helmholtz equation was solved using finite-element discretization with appropriate absorbing or radiation boundary conditions to suppress non-physical inward-propagating waves [44]. Given an estimate of σ(*r*), the electric field *E*_*z*_(*r*) and its squared magnitude |*E*_*z*_(*r*)|^2^ were computed, and the conductivity updating was performed using the relation σ(*r*) = *s*(*r*)/|*E*_*z*_(*r*)|^2^. The forward Helmholtz solution and conductivity update were iterated until the residual error between the DOT-derived power loss density *s*(*r*) and the modeled quantity σ(*r*)|*E*_*z*_(*r*)|^2^ satisfied a predefined convergence criterion. The numerical implementation, finite-element discretization, and regularization strategy follow standard model-based inversion procedures and are conceptually similar to previously reported electrical conductivity reconstruction frameworks in microwave-induced thermoacoustic imaging [45].

### 3.3 Experimental Implementation of OWT

A fast three-dimensional diffuse optical tomography (fast-DOT) system was used for OWT imaging, as schematically illustrated in Fig. 1. The system has been described in detail previously [27, 46]. Briefly, continuous-wave near-infrared light (780 nm) was delivered to the mouse head through source optic fiber bundles, and diffusely scattered light was collected by detection optic fiber bundles. The acquired boundary measurements were digitized and synchronized using dedicated data acquisition hardware under computer control. In this framework, DOT provides the baseline optical measurements from which optical conductivity is subsequently estimated.

**Figure 1.**
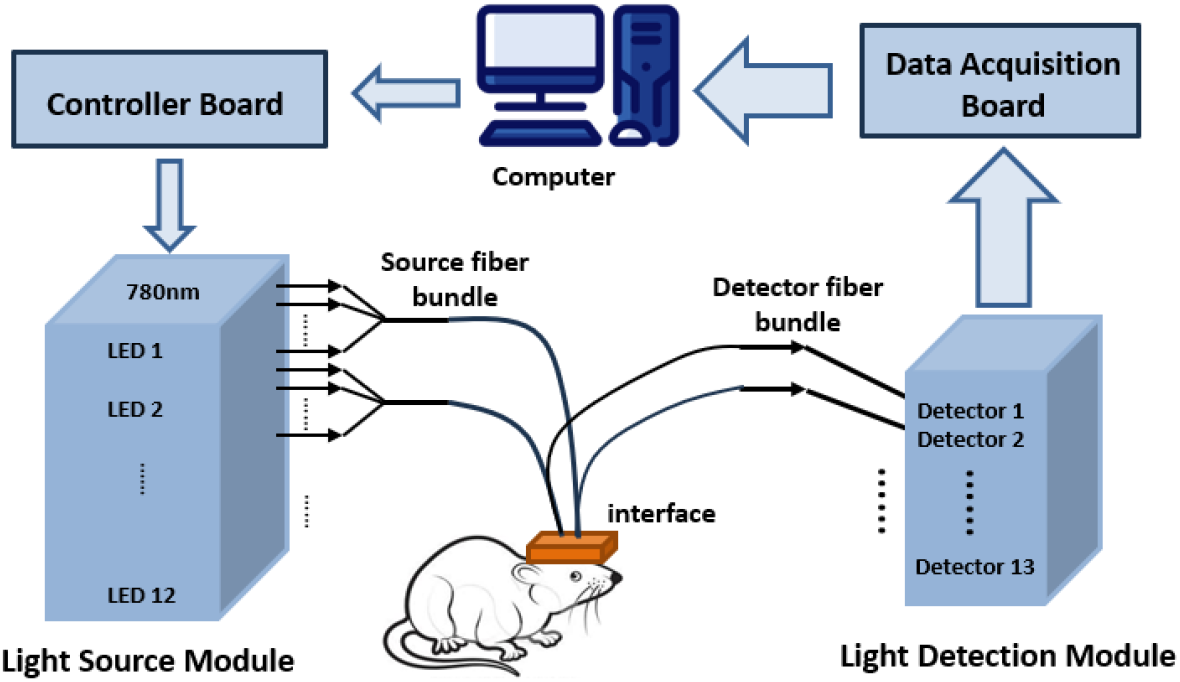
Schematic of our OWT imaging framework. A fast-DOT system for in vivo mouse brain OWT imaging. Continuous-wave near-infrared light (780 nm) is delivered through source optic fiber bundles to the scalp via a miniaturized optical interface, while diffusely scattered light is collected by detection optic fiber bundles and transmitted to the detection system. The acquired signals are digitized and synchronized through data acquisition boards under computer control. The compact interface integrates 12 source optic fiber bundles and 13 detection optic fiber bundles in a dense geometry adapted to the rodent head, enabling non-invasive measurements of light propagation through brain tissue for subsequent image reconstruction.

**Figure 2.**
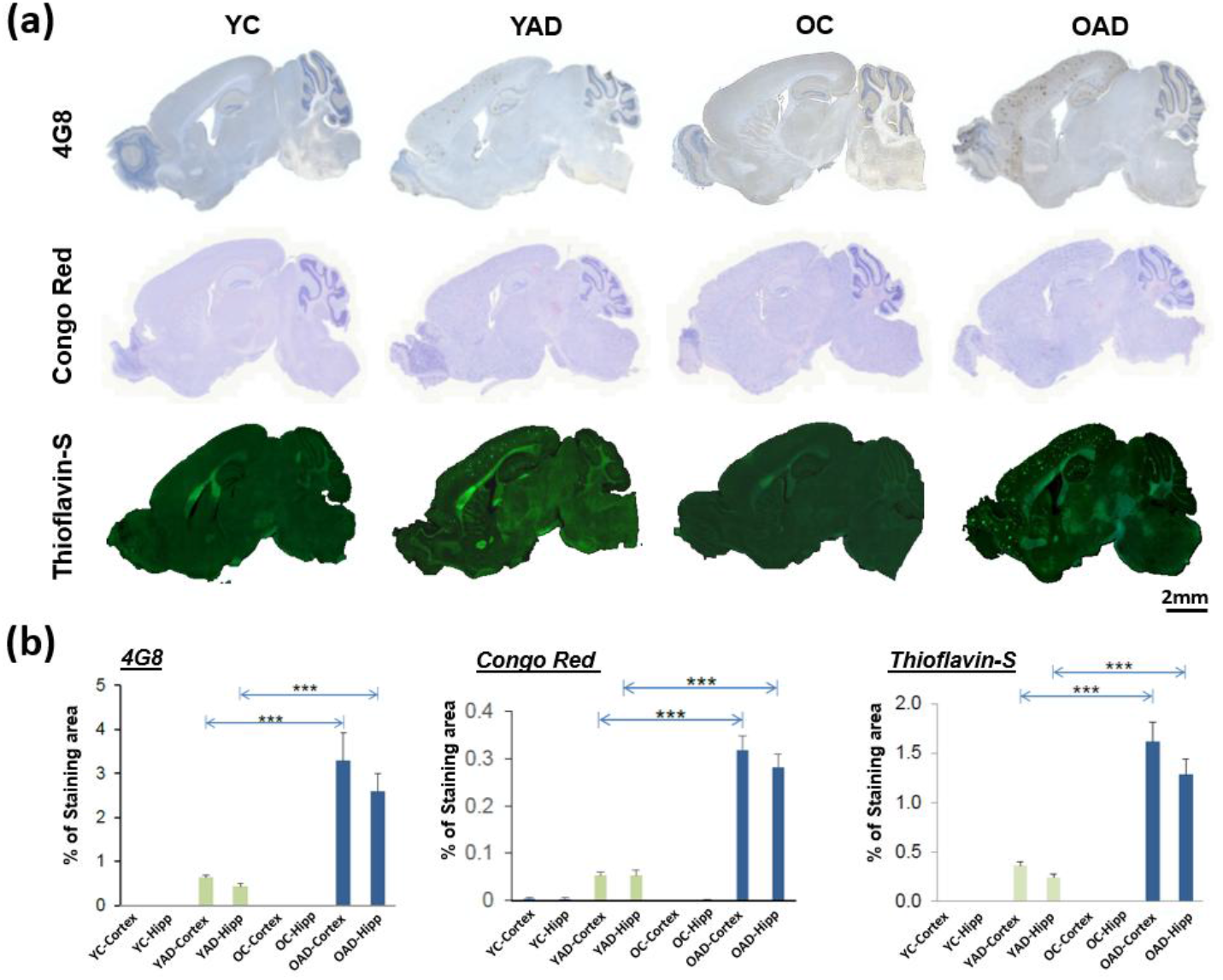
Histological validation of amyloid pathology across age and disease groups. **(a)** Representative sagittal brain sections from YC, YAD, OC, and OAD groups stained with 4G8, Congo Red, and Thioflavin-S. Amyloid deposition increases from control to AD groups and is more extensive in older animals. Scale bar, 2 mm. **(b)** Quantification of amyloid burden (% staining area) in cortex and hippocampus across groups. Data are presented as mean ± s.d.; statistical significance was assessed using two-way ANOVA with post hoc multiple comparisons. ***P < 0.001.

To accommodate the small size and curvature of the mouse head, a miniaturized optical interface was employed to ensure dense spatial sampling and stable optical coupling [46]. The interface integrated 12 source optic fiber bundles and 13 detection optic fiber bundles within a compact geometry (∼17 mm × 17 mm), forming 156 source–detector measurement pairs suitable for in vivo cortical imaging. The measured boundary data were first reconstructed using our finite-element-based inverse model (Eqs. (3)-(5)) to obtain spatial maps of *μ*_*a*_ and 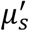. These parameters served as intermediate quantities for subsequent optical conductivity estimation (Eq. (6)).

### 3.4 Animal Model and Experimental Design

An APP/PS1 transgenic mouse model was used to investigate age- and disease-related alterations in optical properties. This model carries human transgenes encoding amyloid precursor protein (APP) and presenilin-1 (PS1), resulting in progressive overproduction and deposition of β-amyloid in the brain. The APP/PS1 line recapitulates key pathological features of Alzheimer’s disease, particularly amyloid plaque formation, and is widely used in preclinical AD research [47-49].

Animals were divided into four experimental groups to evaluate both age and disease effects: young control (YC, ∼7 months, n = 5), young AD (YAD, ∼7 months, n = 6), old control (OC, ∼14.5 months, n = 5), and old AD (OAD, ∼14.5 months, n = 6). Age-matched wild-type littermates served as controls for the transgenic groups. This age-by-disease design was specifically selected to evaluate whether optical conductivity captures stage-dependent alterations, including potential differences between early and later disease states.

For the *in vivo* OWT imaging, the hair over the scalp region was carefully removed to ensure stable optical contact. Animals were anesthetized using inhaled isoflurane gas (4% for induction and 1.5–2% for maintenance) and placed in a stereotaxic frame to minimize motion. The miniaturized optical interface was positioned over the cranial surface, and optical data were recorded continuously for approximately two minutes per animal under steady-state conditions.

### 3.5 Histological Validation

All the *in vivo* OWT imaging experiments were performed prior to tissue collection. Following imaging, animals were euthanized, and brain tissues were harvested for histological validation. Brains were fixed in 10% formalin for 24 hours to preserve tissue architecture, followed by paraffin embedding. Serial coronal sections were prepared using a microtome and mounted onto glass slides for subsequent staining.

To characterize amyloid pathology across disease stages, three complementary staining methods were employed: 4G8 immunostaining, Congo Red, and Thioflavin-S. 4G8 is a monoclonal antibody targeting amyloid-β peptides and enables detection of total amyloid deposition regardless of aggregation state. Congo Red preferentially binds β-sheet–rich fibrillar structures, highlighting mature plaque formations, whereas Thioflavin-S selectively labels highly ordered fibrillar aggregates through fluorescence. The combined use of these staining modalities provides complementary information on plaque presence, distribution, and aggregation state.

Stained sections were examined under bright-field and fluorescence microscopy, and high-resolution images were acquired for quantitative analysis. Plaque-positive areas were measured in predefined cortical and hippocampal regions using standardized image analysis procedures. Histological quantification was performed to establish ground-truth pathological differences across age and disease groups, providing a biological reference for interpreting DOT-derived optical parameters and conductivity measurements. In particular, this analysis enables evaluation of whether imaging-derived changes correspond to early versus advanced amyloid pathology. All the procedures were conducted in accordance with institutional animal care guidelines and approved protocols.

### 3.6 Statistical Analysis

Descriptive statistics were calculated for each optical parameter (*μ*_*a*_, 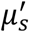, and σ) and are reported as mean ± s.d. for each age and disease group. To evaluate the independent and combined effects of age (young vs. old) and disease status (control vs. AD), two-way analysis of variance (ANOVA) was performed with interaction terms included [50]. When appropriate, post hoc pairwise comparisons were conducted with correction for multiple testing [51]. Effect sizes were quantified using partial *η*^2^ for ANOVA and Cohen’s d for pairwise comparisons [52].

The primary statistical objective was to determine whether optical conductivity exhibits an age-by-disease interaction indicative of stage-dependent effects, in contrast to conventional optical parameters. To assess diagnostic performance, logistic regression models were constructed [53] using leave-one-out cross-validation (LOOCV) [54]. For each fold, features were standardized using training data only, and the trained model was applied to the held-out subject to obtain an out-of-fold predicted probability. Receiver operating characteristic (ROC) curves were generated from these probabilities, and the area under the curve (AUC) was calculated to quantify discriminative ability [55]. Classification analyses were performed separately within each age cohort to evaluate stage-specific diagnostic performance, particularly for early-stage disease. Both individual optical parameters and their combinations were assessed to determine whether optical conductivity provides complementary information beyond conventional metrics. All statistical analyses were performed in MATLAB (MathWorks, Natick, MA).

## 4. Results and Discussion

Before presenting and interpreting the in vivo optical imaging results, we attempt to establish pathological ground truth using the ex vivo histological results, as shown in Fig.2. The three staining modalities show consistent agreement in both the spatial distribution and relative extent of amyloid deposition, enabling reliable comparison of plaque burden across age and disease groups. Control groups (YC and OC) exhibit minimal staining, whereas YAD shows early-stage plaque emergence, and OAD demonstrates a pronounced increase in amyloid deposition across both cortex and hippocampus. Quantitative analysis further confirms a significant elevation of plaque burden in OAD relative to all other groups (p < 0.001), with YAD exhibiting intermediate levels and OC remaining comparable to YC.

These histological results establish a clear disease-driven, stage-dependent progression of amyloid pathology that is not attributable to aging alone. This well-defined pathological gradient provides a biologically grounded reference for assessing whether imaging-derived optical parameters can capture both the presence of pathology and its progression across disease stages.

### 4.1 Image of Optical Parameters Across Age and Disease Groups

Figure 3 presents the spatial distributions of *μ*_*a*_, 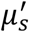, and σ with anatomical and depth context for representative mouse brains. Panel (a) reveals that σ varies remarkably along the dorsal–ventral axis, with localized regions of increased values emerging at different depths, consistent with areas of elevated pathological burden, indicating depth-dependent heterogeneity rather than a uniform volumetric distribution. In panel (b), the *μ*_*a*_ maps remain relatively uniform across both cortical and subcortical areas, including the hippocampal region, showing limited spatial variation. In contrast, 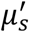 exhibits a smoother but elevated distribution in AD groups, with increased values broadly distributed across the cortex and extending into deeper structures. More notably, σ displays a distinct spatial organization. Regions of elevated σ appear as heterogeneous and localized structures, with prominent increases observed in cortical regions and portions of the hippocampal area, particularly in AD groups, broadly consistent with areas of elevated pathological burden. These high σ regions are not strictly co-localized with the dominant gradients of *μ*_*a*_ or 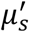, and in some cases emerge in regions where neither *μ*_*a*_ nor 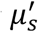 shows pronounced local maxima. Together, these observations indicate that σ exhibits structured, spatially heterogeneous variations that are not detected by *μ*_*a*_ or 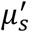, suggesting that it captures additional tissue characteristics beyond conventional optical parameters and motivating subsequent quantitative evaluation.

**Figure 3.**
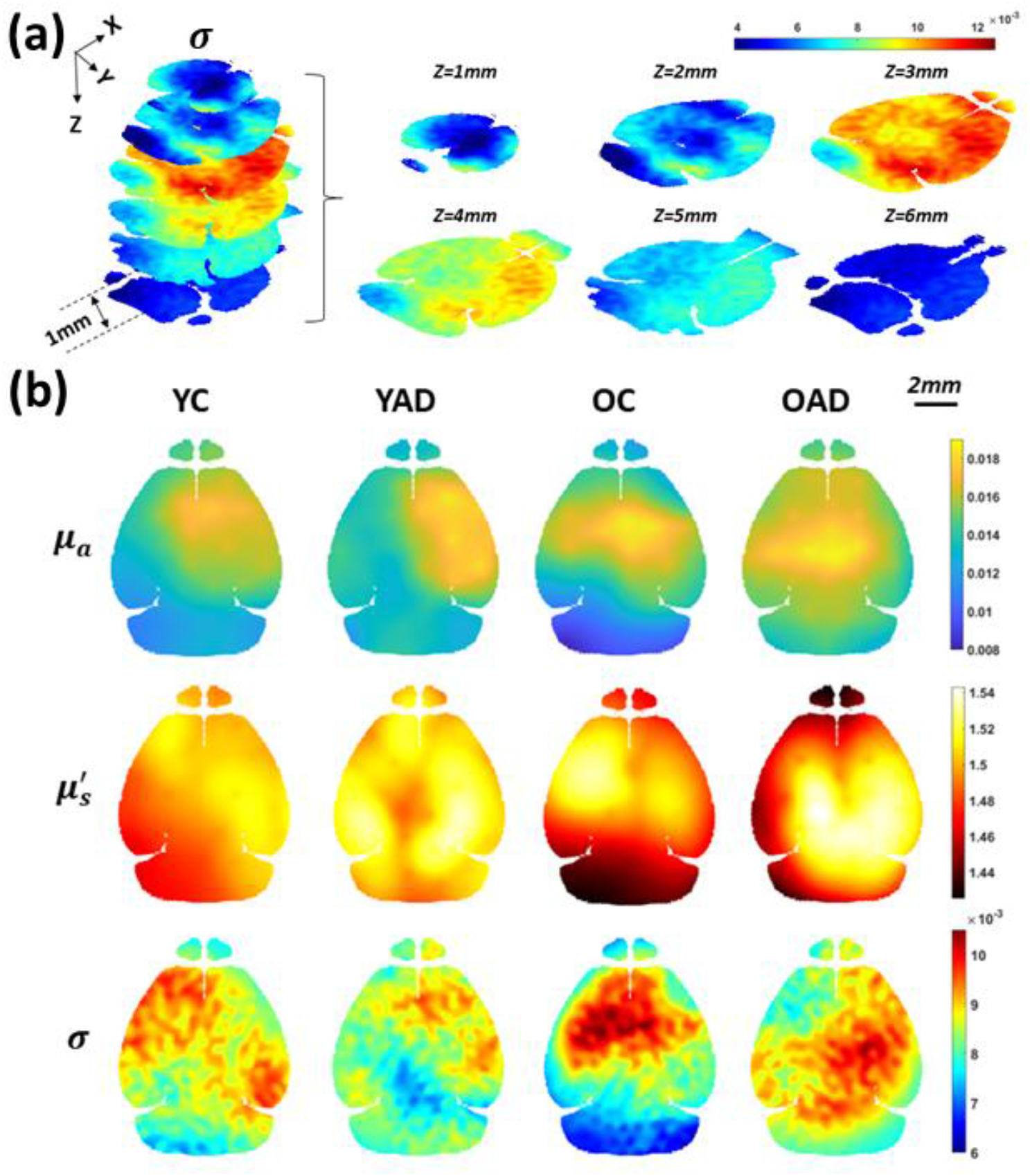
Representative spatial distributions of optical parameters across age and disease groups. (a) Three-dimensional rendering of σ image for a representative YAD mouse brain. The first column shows a stacked visualization across multiple axial (transverse) slices, illustrating volumetric distribution of σ. The second to the fourth columns present corresponding individual axial slices sampled at 1 mm intervals from the dorsal surface, z = 1–6 mm where z denotes depth along the dorsal–ventral axis as defined in the Allen Mouse Brain Atlas. (b) Representative *μ*_*a*_, 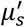 and σ images across all groups, shown as an axial (transverse) slice at a fixed depth (z = 3 mm) for direct comparison. Color bars (top right for (a) and right for (b)) represent the quantitative values of *μ*_*a*_(mm^-1^), 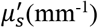 and σ(S/m), respectively. Brain outlines are derived from the Allen Mouse Brain Atlas. Scale bar, 2 mm.

To determine whether the spatially heterogeneous distributions observed in Fig. 3 correspond to consistent and quantifiable group-level differences, we examined the quantitative distributions of *μ*_*a*_, 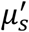, and σ across age and disease groups (Fig. 4; summary statistics in Table S1). *μ*_*a*_ values are highly similar across all groups (Fig. 4a), with mean values ranging from approximately 0.018 to 0.019 mm^−1^ and strongly overlapping distributions. The corresponding violin plots show narrow spreads and comparable shapes across groups, with no evident shifts in central tendency or dispersion, indicating minimal sensitivity of *μ*_*a*_ to either age or disease. In contrast, 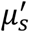 values demonstrate a clear disease-related increase (Fig. 4b). Mean 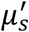 values are elevated in AD animals compared to age-matched controls (YC: 1.99 ± 0.17 mm^−1^ vs. YAD: 2.32 ± 0.31 mm^−1^; OC: 1.92 ± 0.10 mm^−1^ vs. OAD: 2.44 ± 0.47 mm^−1^), corresponding to an approximate 15–25% increase in scattering for AD groups. These differences in scattering are also reflected in the distribution shapes, where AD groups show a noticeable shift toward higher values accompanied by broader spread and reduced overlap with controls. This pattern is consistent across age cohorts, suggesting that 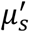 captures a generalized disease-associated shift compared to controls in tissue microstructure. σ values are noted to exhibit a qualitatively distinct behavior differing from both *μ*_*a*_ and 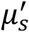 (Fig. 4c). In the young cohort, σ is reduced in AD animals relative to controls (YC: 0.023 ± 0.002 S/m vs. YAD: 0.020 ± 0.001 S/m; ∼13% decrease), which is reflected in a shift of the distribution toward lower values with relatively tight spread. In the old cohort, this pattern is attenuated or partially reversed, with OAD animals showing slightly higher σ values than OC animals (OAD: 0.022 ± 0.001 S/m vs. OC: 0.021 ± 0.002 S/m). Notably, the distributions in the older groups exhibit greater variability and overlap, compared to the young cohort.

**Figure 4.**
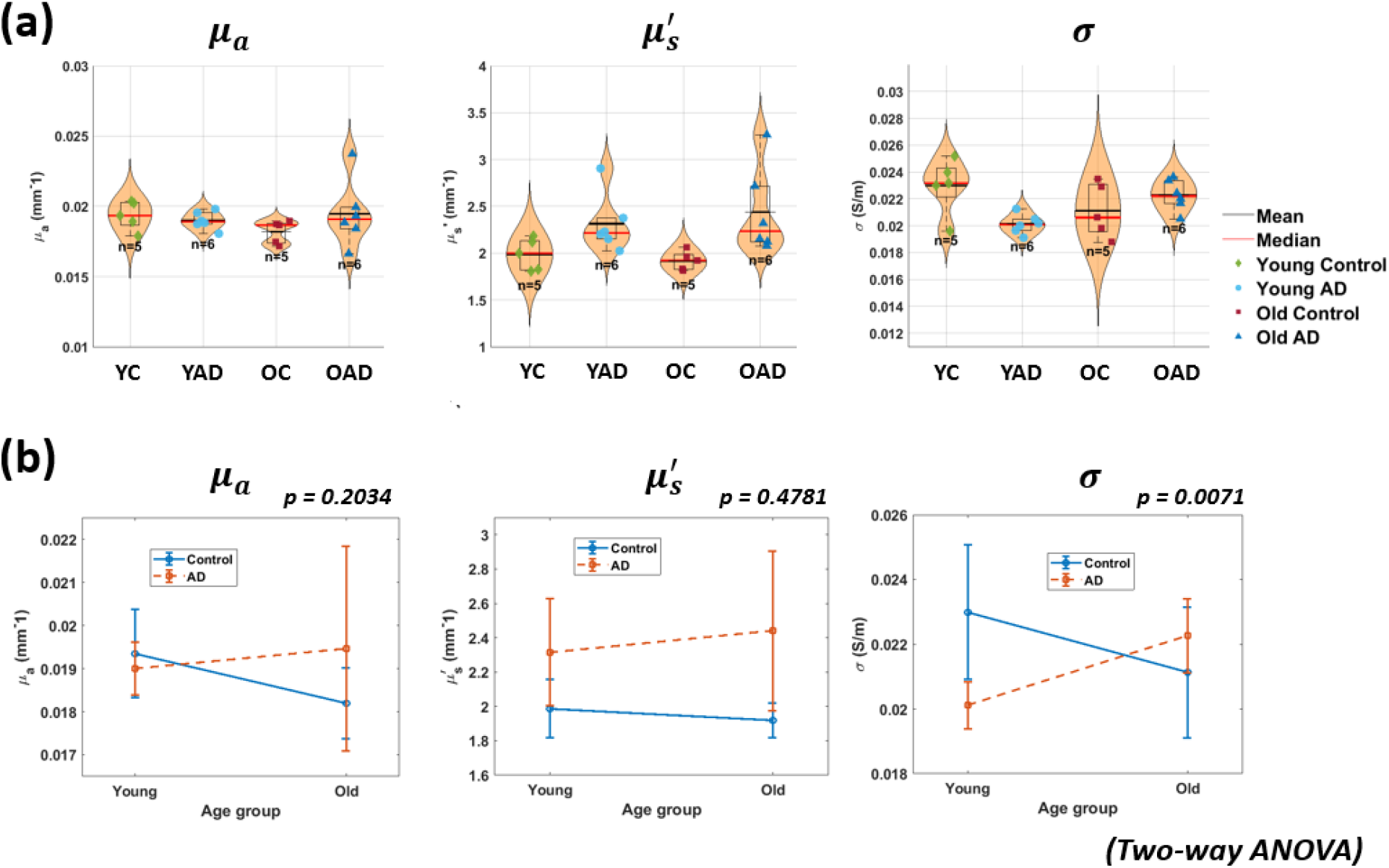
Quantitative distributions and age-by-disease interaction of optical parameters. (a) Violin plots show the distributions of *μ*_*a*_(left), 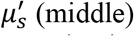, and σ (right) across all groups. These quantitative values were extracted from their respective images for each mouse, corresponding to the sampling depth determined by the optical interface configuration. This depth minimizes surface-related artifacts while encompassing major brain regions, including cortex and hippocampus, and provides a consistent basis for comparison without relying on precise spatial localization. Individual data points are overlaid, with mean (black) and median (red) indicated. (b) Interaction plots summarize age-by-disease effects for each parameter, showing group means for control and AD conditions across young and old cohorts. Absorption and scattering exhibit no significant interaction, whereas optical conductivity shows a clear crossover pattern, consistent with a significant age × disease interaction (two-way ANOVA, p=0.0071). Quantitative values were defined as the maximum reconstructed parameter within a depth range of 2–6 mm for all panels.

Together, these findings indicate that while conventional optical parameters primarily reflect either minimal (for *μ*_*a*_) or generalized (for 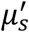) disease effects, σ exhibits age-dependent directional changes that reflect an interaction-driven effect, which are not observed in *μ*_*a*_ or 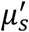. These changes are further accompanied by differences in distribution shape and variability, indicating that σ captures a distinct mode of tissue alteration consistent with dynamic, stage-dependent microenvironmental reorganization.

### 4.2 Conventional Optical Parameters (*μ*_*a*_ and 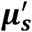) Lack Stage-Specific Sensitivity

We now statistically evaluate the performance of DOT-derived conventional optical parameters (*i. e*., *μ*_***a***_ and 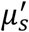) in terms of disease sensitivity and stage differentiation. As shown in Figure 4(a) and Table S1, *μ*_***a***_ values are highly similar across all groups, with nearly identical means (≈0.018–0.019 mm^−1^) and substantial overlap in distributions. Two-way ANOVA reveals no significant main effects of age or disease and no age-by-disease interaction (all p > 0.20; Table S2). Pairwise comparisons are likewise non-significant (Table S3), indicating minimal sensitivity of *μ*_***a***_ to AD-related pathology in this model. In contrast, 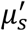 values demonstrate a consistent disease-related increase across both age cohorts. As shown in Figure 4(a), mean 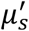 values were elevated in AD animals compared to age-matched controls (YC: 1.99 ± 0.17 mm^−1^vs. YAD: 2.32 ± 0.31 mm^−1^; OC: 1.92 ± 0.10 mm^−1^vs. OAD: 2.44 ± 0.47 mm^−1^), corresponding to an approximate 15–25% increase. This pattern is confirmed by a significant main effect of disease (F = 10.26, p = 0.0049; Table S2), while neither the main effect of age nor the age-by-disease interaction reaches significance.

These findings indicate that 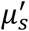 captures a generalized disease-associated change in tissue microstructure but does not differentiate between early and later disease stages. Together, conventional optical parameters exhibit either minimal sensitivity (*μ*_***a***_) or non-stage-specific sensitivity 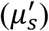, highlighting the need for complementary imaging metrics.

### 4.3 Optical Conductivity Reveals an Age-Dependent Disease Interaction

In contrast to conventional optical parameters, σ demonstrates a pronounced age-dependent disease interaction. As shown in Figure 4(a) and Table S1, in the young group, AD animals show reduced conductivity relative to controls (YC: 0.023 ± 0.002 S/m vs. YAD: 0.020 ± 0.001 S/m; ∼13% decrease). In the old cohort, such relationship is changed, with OAD animals demonstrating slightly higher σ values than OC animals (OAD: 0.022 ± 0.001 S/m vs. OC: 0.021 ± 0.002 S/m). This crossover pattern is statistically confirmed by two-way ANOVA, which reveals a significant age-by-disease interaction (F = 9.20, p = 0.0071; Fig. 4b; full statistics in supplementary Table S2), while neither the main effect of age nor disease reaches significance. The presence of a significant interaction in the absence of dominant main effects indicates that the impact of AD on σ is fundamentally stage-dependent rather than uniform across disease progression.

Pairwise comparisons further support this interpretation. In the young cohort, σ values are significantly reduced in AD animals relative to controls (p = 0.034, Cohen’s d = 1.93; Table S3), indicating a large effect size. In contrast, no significant difference is observed between OAD and OC animals. Importantly, this opposing direction of change across age groups would be obscured in pooled analyses, explaining the absence of a main disease effect. Thus, σ uniquely captures a stage-dependent disease response that is not detectable using conventional optical parameters.

### 4.4 Early-Stage Sensitivity of Optical Conductivity

In the young cohort, σ achieves an AUC of 0.80, compared with 0.27 for *μ*_***a***_ and 0.73 for 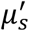 (Table 1). This strong classification performance is consistent with the large effect size observed in the young cohort (Cohen’s d = 1.93), indicating that conductivity changes are both statistically robust and diagnostically meaningful at early disease stages.

**Table 1.** Diagnostic performance (AUC) of individual and combined optical parameters. Classification performance evaluated using LOOCV logistic regression is summarized for individual optical parameters (*μ*_*a*_, 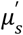, σ) and their combinations. Diagnostic performance is reported as the area under the receiver operating characteristic curve (AUC). Classification was performed based on the quantitative values defined in Figure 4.

| Model | Young cohort AUC | Old cohort AUC |
| --- | --- | --- |
| $\mu_a$ | 0.27 | 0.43 |
| $\mu'_s$ | 0.73 | **0.87** |
| $\sigma$ | 0.80 | 0.50 |
| $\sigma + \mu'_s$ | 0.83 | **0.87** |
| $\sigma + \mu'_s + \mu_a$ | **0.87** | 0.80 |

In contrast, σ demonstrates limited discriminative performance in the old cohort (AUC = 0.50), consistent with the absence of significant group differences between OAD and OC animals. Although 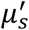 achieved higher AUC values in the old cohort (AUC = 0.87), this reflects a generalized disease-related alteration rather than stage-specific modulation. These findings indicate that σ is particularly sensitive to early-stage pathological changes, capturing alterations that are not fully reflected in conventional optical parameters. The concordance between effect size and classification performance further supports that σ reflects biologically meaningful changes rather than statistical variability.

### 4.5 Multimodal Combination Enhances Discrimination in the Young Cohort

Finally, we evaluate whether combining optical parameters improves diagnostic performance, particularly in the young cohort where σ exhibits strong stage-dependent sensitivity. As shown in Table 1 and Supplementary Figure S1, combining σ with 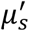 increases classification performance in the young cohort (AUC = 0.83), while integration of all three parameters (*μ*_***a***_, 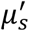, σ) further improves AUC to 0.87. These results indicate that σ provides complementary information beyond conventional optical metrics.

Notably, this improvement is not observed in the old cohort, where multimodal models do not substantially outperform 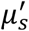 alone. This difference is consistent with the interaction-driven behavior of σ: in early-stage disease, σ captures additional biological variation not reflected in scattering, whereas in later stages, scattering-related changes dominate. Together, these findings demonstrate that multimodal integration enhances early-stage discrimination, with σ serving as a critical contributor to improved diagnostic performance.

### 4.6 Mechanistic Insights into Conductivity Alterations

Optical conductivity characterizes tissue response to optical-frequency electromagnetic waves and reflects integrated energy dissipation shaped by absorption, dielectric properties, and microstructure-dependent field redistribution. Unlike *μ*_***a***_, which primarily reflects chromophore concentration, and 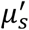, which captures bulk scattering from refractive index heterogeneity, σ represents a wave/field-weighted measure of how electromagnetic energy is distributed and dissipated within tissue.

The observed age-dependent interaction suggests that σ is sensitive to dynamic microenvironmental changes that evolve across disease stages. The reduced optical conductivity observed in young AD animals likely reflects early alterations in tissue organization, such as synaptic loss, subtle neuronal degeneration, or changes in membrane structure and molecular packing, which modify dielectric properties without producing large scattering changes. As disease progresses, secondary processes including gliosis, extracellular matrix remodeling, and changes in water content may alter electromagnetic energy dissipation pathways, contributing to the attenuation or partial reversal of σ differences observed in older animals. This crossover behavior indicates that σ does not simply track cumulative pathology but instead reflects evolving tissue reorganization. Importantly, because σ is derived through a coupled electromagnetic wave model rather than a simple algebraic combination of *μ*_***a***_ and 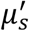, it captures nontrivial interactions between absorption and scattering that are not accessible through conventional optical parameters alone. This integrative sensitivity provides a mechanistic basis for the enhanced early-stage discrimination observed for σ and supports its potential as a complementary biomarker for characterizing neurodegenerative processes.

### 4.7 Limitations

Several limitations exist in this study. First, the sample size within each group was modest, which may limit statistical power and the generalizability of the observed effects. While consistent trends and large effect sizes were observed, particularly for optical conductivity in the young cohort, larger size studies are needed to confirm the robustness and reproducibility of these results. Second, this study was conducted in a transgenic mouse model of Alzheimer’s disease. While the APP/PS1 model captures key features of amyloid pathology, it does not fully reflect the complexity of human disease, including tau-related neurodegeneration and clinical heterogeneity. Therefore, translation of these findings to human populations will require further validation. Third, optical conductivity is a new imaging parameter derived from OWT framework. Although grounded in established electromagnetic and diffuse optical principles, its biological interpretation remains indirect. The observed associations with microstructural remodeling are therefore inferential and will benefit from further validation using multimodal approaches, including histological, electrophysiological, or complementary imaging techniques. Fourth, the estimation of optical conductivity depends on intermediate reconstruction steps, including absorption, scattering, and photon fluence, and electric field. Uncertainty or error in these intermediate quantities may propagate into conductivity estimates. Future work should evaluate the sensitivity of the reconstruction pipeline to measurement noise, model assumptions, and parameter initialization. Finally, the present study uses a cross-sectional design across two age groups, limiting the ability to directly assess longitudinal changes within individual animals. Longitudinal studies will be necessary to determine how optical conductivity evolves over time and how it relates to disease progression and functional outcomes.

## 5. Conclusions

In this work, we have introduced optical wave tomography as a novel imaging framework for reconstructing optical conductivity and demonstrated its utility in characterizing stage-dependent alterations in an Alzheimer’s disease model. By formulating optical conductivity as a power-loss–based parameter derived from diffuse optical measurements and estimating it through a coupled electromagnetic wave model, OWT establishes a new approach for quantifying tissue electromagnetic energy dissipation in vivo. Unlike conventional optical parameters, which exhibited either minimal sensitivity (absorption) or generalized, non-stage-specific changes (scattering), optical conductivity revealed a significant age-by-disease interaction and a distinct stage-dependent crossover pattern. This finding indicates that optical conductivity captures dynamic microenvironmental remodeling that evolves across disease progression, rather than simply reflecting cumulative pathological burden. OWT addresses a key limitation in current Alzheimer’s disease imaging strategies, i.e., the inability to detect and characterize intermediate tissue changes that link early molecular pathology to later structural degeneration. The observed early-stage sensitivity of optical conductivity, supported by both effect size and classification performance, suggests that OWT provides complementary information beyond existing imaging approaches. More broadly, optical conductivity represents a shift from conventional optical imaging parameters toward an integrated, optical wave-based description of tissue organization, offering a new perspective on the relationship between microstructure and function in neurodegenerative disease. In sum, OWT establishes optical conductivity as a methodologically innovative and biologically informative imaging biomarker with potential relevance for early-stage detection, disease staging, and mechanistic investigation of Alzheimer’s disease and related neurodegenerative disorders.

## Contributions

**H.Y**. designed and performed the experiments, analyzed the data, and wrote the manuscript. **S.J**. provided expertise in Alzheimer’s disease pathology and wrote the manuscript. **H.J**. conceived and supervised the study, and wrote the manuscript.

## Supplementary Material

**Figure S1.**
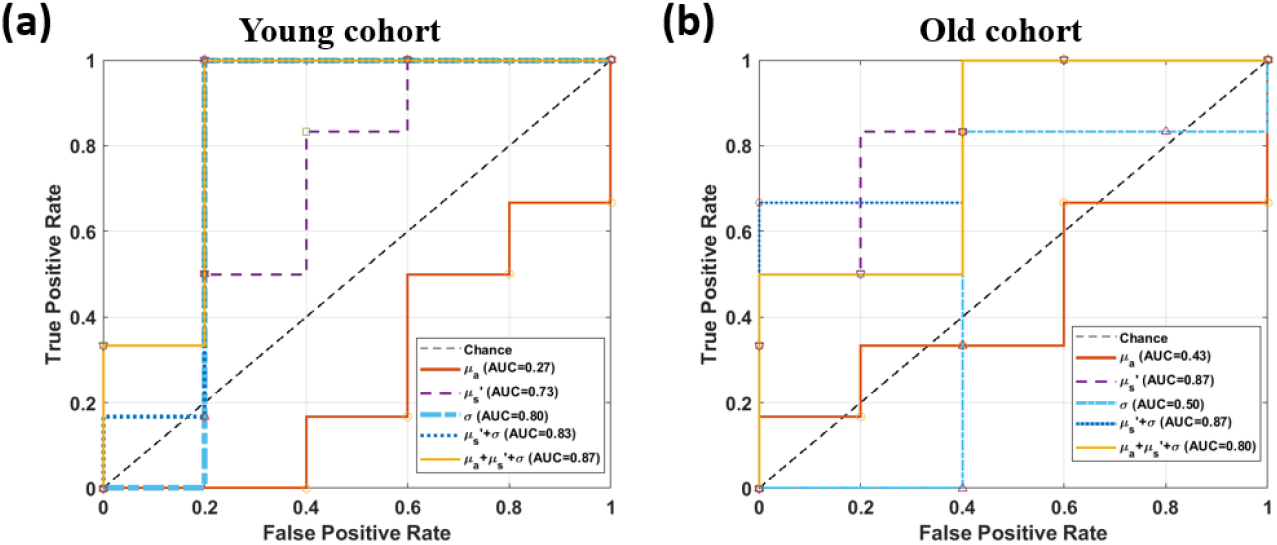
ROC curves of individual and combined optical parameters by age cohort. ROC curves for classification of control and AD groups using individual optical parameters (*μ*_*a*_, 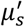, σ) and their combinations. Results are shown for **(a)** the young cohort and **(b)** the old cohort. Classification performance was evaluated using LOOCV logistic regression, with corresponding AUC values summarized in Table 1.

**Table S1.**
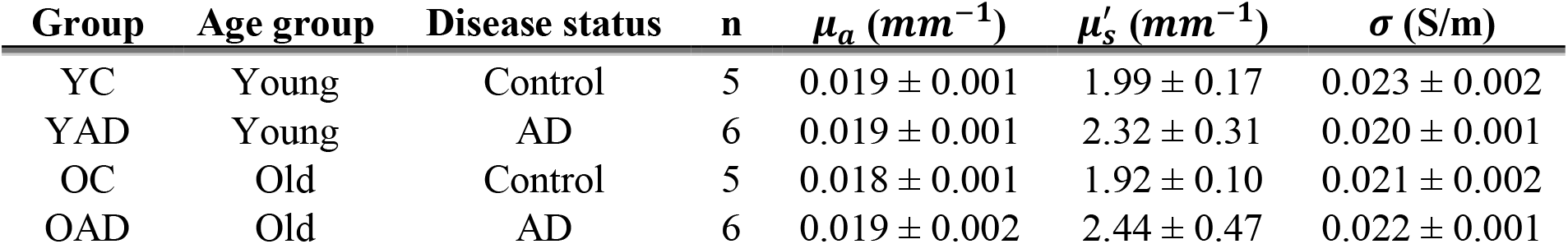
Summary statistics of optical parameters by age group and disease status. Mean values (± s.d.) of *μ*_*a*_, 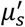, and σ are reported for each group (YC, YAD, OC, OAD), where quantitative values for each mouse were extracted and calculated as described in Figure 4. Two-way ANOVA was performed to assess the main effects of age and disease, as well as their interaction, with corresponding p-values reported.

**Table S2.** Two-way ANOVA results for age, disease, and age-by-disease interaction effects. Two-way ANOVA results for *μ*_***a***_, 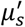, and σ are shown, evaluating the main effects of age and disease, as well as their interaction (age × disease). For each outcome, F-statistics and corresponding p-values are reported. Statistically significant effects (p < 0.05) are highlighted in bold. Quantitative values were extracted and calculated as described in Figure 4.

| Outcome | Effect | df | F | p-value |
| --- | --- | --- | --- | --- |
| $\mu_a$ | Age | 1.18 | 0.19 | 0.6655 |
|  | Disease | 1.18 | 0.56 | 0.4643 |
| | Age × Disease | 1.18 | 1.74 | 0.2034 |
| $\mu'_s$ | Age | 1.18 | 0.080 | 0.7800 |
|  | **Disease** | 1.18 | **10.26** | **0.0049** |
| | Age × Disease | 1.18 | 0.53 | 0.4781 |
| $\sigma$ | Age | 1.18 | 0.25 | 0.6253 |
|  | Disease | 1.18 | 1.73 | 0.2044 |
|  | **Age × Disease** | 1.18 | **9.20** | **0.0071** |

**Table S3.**
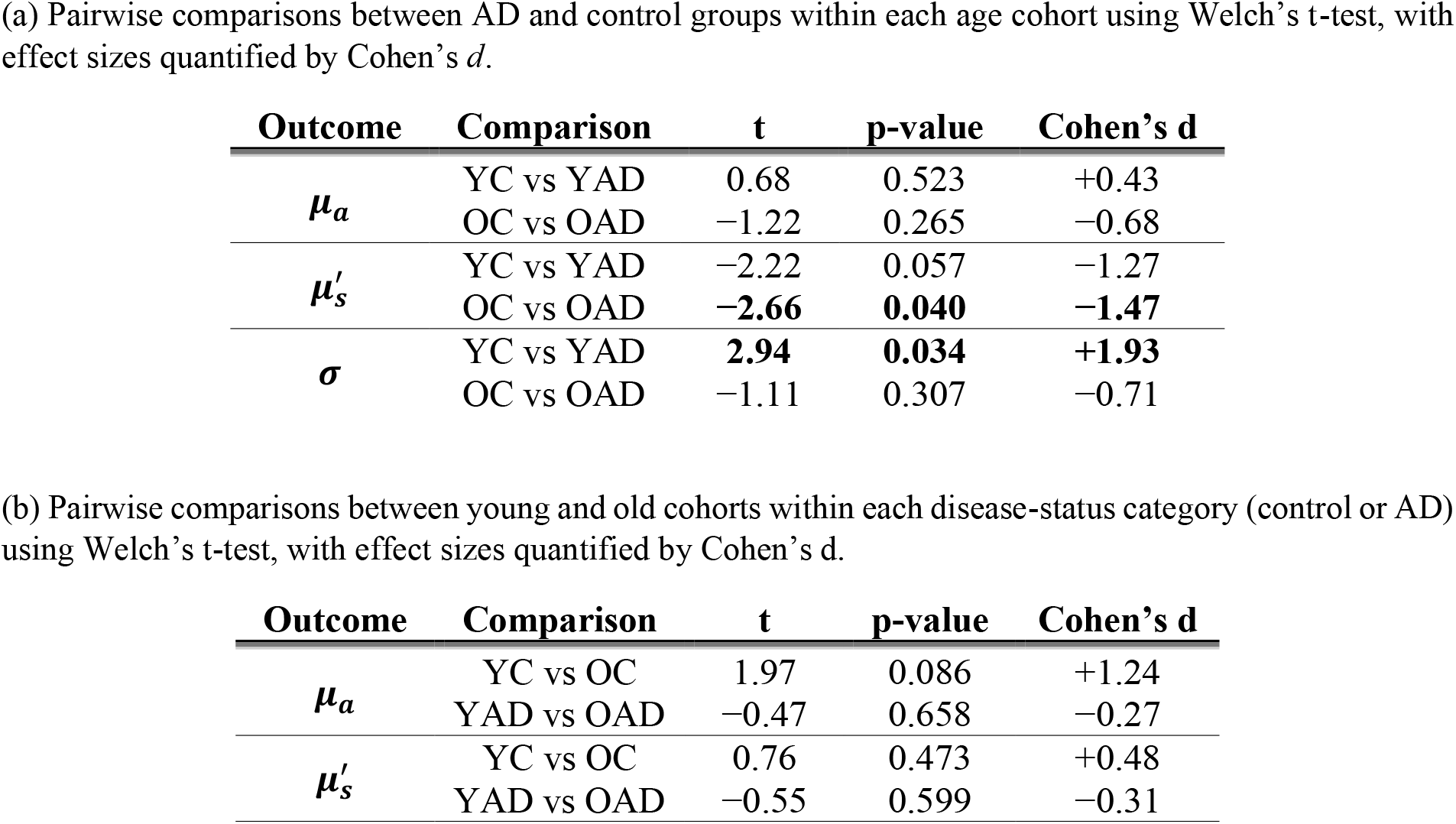
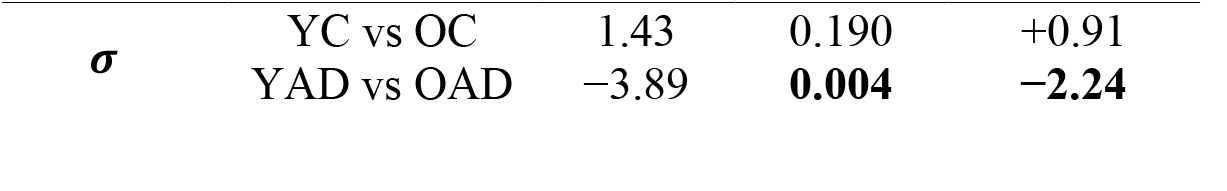
Additional statistical analyses and pairwise group comparisons.

## References

1. Nichols, E., et al., Estimation of the global prevalence of dementia in 2019 and forecasted prevalence in 2050: an analysis for the Global Burden of Disease Study 2019. The Lancet Public Health, 2022. 7(2): p. e105–e125.

2. Prince, M., et al., The global prevalence of dementia: a systematic review and metaanalysis. Alzheimer’s & dementia, 2013. 9(1): p. 63–75. e2.

3. DeTure, M.A. and D.W. Dickson, The neuropathological diagnosis of Alzheimer’s disease. Molecular neurodegeneration, 2019. 14(1): p. 32.

4. Braak, H. and E. Braak, Neuropathological stageing of Alzheimer-related changes. Acta neuropathologica, 1991. 82(4): p. 239–259.

5. Selkoe, D.J. and J. Hardy, The amyloid hypothesis of Alzheimer’s disease at 25 years. EMBO molecular medicine, 2016. 8(6): p. 595–608.

6. Jack, C.R., et al., Hypothetical model of dynamic biomarkers of the Alzheimer’s pathological cascade. The Lancet Neurology, 2010. 9(1): p. 119–128.

7. Sperling, R.A., et al., Toward defining the preclinical stages of Alzheimer’s disease: Recommendations from the National Institute on Aging-Alzheimer’s Association workgroups on diagnostic guidelines for Alzheimer’s disease. Alzheimer’s & dementia, 2011. 7(3): p. 280–292.

8. Jellinger, K., Towards a biological definition of Alzheimer disease. Int J Neurol Neurother, 2020. 7(1): p. 095.

9. Hampel, H., et al., Blood-based biomarkers for Alzheimer disease: mapping the road to the clinic. Nature Reviews Neurology, 2018. 14(11): p. 639–652.

10. McKhann, G., et al., The diagnosis of dementia due to Alzheimer’s disease: Recommendations from the NIAA Association workgroups on diagnostic guidelines for AD. Alzheimers Dement, 2011. 7(3): p. 263–269.

11. Dubois, B., et al., Advancing research diagnostic criteria for Alzheimer’s disease: the IWG-2 criteria. The Lancet Neurology, 2014. 13(6): p. 614–629.

12. Ossenkoppele, R., et al., Tau PET patterns mirror clinical and neuroanatomical variability in Alzheimer’s disease. Brain, 2016. 139(5): p. 1551–1567.

13. Leuzy, A., et al., Tau PET imaging in neurodegenerative tauopathies—still a challenge. Molecular psychiatry, 2019. 24(8): p. 1112–1134.

14. Saint-Aubert, L., et al., Tau PET imaging: present and future directions. Molecular neurodegeneration, 2017. 12(1): p. 19.

15. Blennow, K., et al., Cerebrospinal fluid and plasma biomarkers in Alzheimer disease. Nature Reviews Neurology, 2010. 6(3): p. 131–144.

16. Karikari, T.K., et al., Blood phosphorylated tau 181 as a biomarker for Alzheimer’s disease: a diagnostic performance and prediction modelling study using data from four prospective cohorts. The Lancet Neurology, 2020. 19(5): p. 422–433.

17. Janelidze, S., et al., Plasma P-tau181 in Alzheimer’s disease: relationship to other biomarkers, differential diagnosis, neuropathology and longitudinal progression to Alzheimer’s dementia. Nature medicine, 2020. 26(3): p. 379–386.

18. Mattsson, N., et al., Plasma tau in Alzheimer disease. Neurology, 2016. 87(17): p. 1827–1835.

19. Jack Jr, C.R., et al., A/T/N: An unbiased descriptive classification scheme for Alzheimer disease biomarkers. Neurology, 2016. 87(5): p. 539–547.

20. Jagust, W.J., et al., Relationships between biomarkers in aging and dementia. Neurology, 2009. 73(15): p. 1193–1199.

21. Mattsson-Carlgren, N., et al., The implications of different approaches to define AT (N) in Alzheimer disease. Neurology, 2020. 94(21): p. e2233–e2244.

22. Ferrari, M. and V. Quaresima, A brief review on the history of human functional near-infrared spectroscopy (fNIRS) development and fields of application. Neuroimage, 2012. 63(2): p. 921–935.

23. Boas, D.A., et al., Twenty years of functional near-infrared spectroscopy: introduction for the special issue. 2014, Elsevier. p. 1–5.

24. Lin, A.J., et al., Optical imaging in an Alzheimer’s mouse model reveals amyloid-β-dependent vascular impairment. Neurophotonics, 2014. 1(1): p. 011005–011005.

25. Lin, A.J., et al., In vivo optical signatures of neuronal death in a mouse model of Alzheimer’s disease. Lasers in surgery and medicine, 2014. 46(1): p. 27–33.

26. Dai, X., et al., Fast noninvasive functional diffuse optical tomography for brain imaging. Journal of biophotonics, 2018. 11(3): p. e201600267.

27. Yang, H., et al., In vivo imaging of epileptic foci in rats using a miniature probe integrating diffuse optical tomography and electroencephalographic source localization. Epilepsia, 2015. 56(1): p. 94–100.

28. Tromberg, B.J., et al., Assessing the future of diffuse optical imaging technologies for breast cancer management. Medical physics, 2008. 35(6Part1): p. 2443–2451.

29. Jiang, H., Diffuse optical tomography: principles and applications. 2018: CRC press.

30. De Strooper, B. and E. Karran, The cellular phase of Alzheimer’s disease. Cell, 2016. 164(4): p. 603–615.

31. Weston, P.S., et al., Diffusion imaging changes in grey matter in Alzheimer’s disease: a potential marker of early neurodegeneration. Alzheimer’s research & therapy, 2015. 7(1): p. 47.

32. Syková, E. and C. Nicholson, Diffusion in brain extracellular space. Physiological reviews, 2008. 88(4): p. 1277–1340.

33. Park, S., et al., Application of high-frequency conductivity map using MRI to evaluate it in the brain of Alzheimer’s disease patients. Frontiers in Neurology, 2022. 13: p. 872878.

34. Hoy, A.R., et al., Microstructural white matter alterations in preclinical Alzheimer’s disease detected using free water elimination diffusion tensor imaging. PloS one, 2017. 12(3): p. e0173982.

35. Jackson, J.D. and R.F. Fox, Classical electrodynamics. 1999, American Association of Physics Teachers.

36. Landau, L.D., et al., Electrodynamics of continuous media. Vol. 8. 2013: elsevier.

37. Stratton, J.A., Electromagnetic theory. 2007: John Wiley & Sons.

38. Ishimaru, A., Wave propagation and scattering in random media. Vol. 2. 1978: Academic press New York.

39. Arridge, S.R., Optical tomography in medical imaging. Inverse problems, 1999. 15(2): p. R41–R93.

40. Jacques, S.L., Optical properties of biological tissues: a review. Physics in medicine and biology, 2013. 58(11): p. R37–R61.

41. Cheong, W.-F., S.A. Prahl, and A.J. Welch, A review of the optical properties of biological tissues. IEEE journal of quantum electronics, 2002. 26(12): p. 2166–2185.

42. Gabriel, C., S. Gabriel, and E. Corthout, The dielectric properties of biological tissues: I. Literature survey. Physics in medicine & biology, 1996. 41(11): p. 2231–2249.

43. Schwan, H.P., Electrical properties of tissue and cell suspensions, in Advances in biological and medical physics. 1957, Elsevier. p. 147–209.

44. Jin, J.-M., The finite element method in electromagnetics. 2015: John Wiley & Sons.

45. Yao, L., G. Guo, and H. Jiang, Quantitative microwave - induced thermoacoustic tomography. Medical physics, 2010. 37(7Part1): p. 3752–3759.

46. Zhang, T., et al., Pre-seizure state identified by diffuse optical tomography. Scientific reports, 2014. 4(1): p. 3798.

47. Jankowsky, J.L., et al., Co-expression of multiple transgenes in mouse CNS: a comparison of strategies. Biomolecular engineering, 2001. 17(6): p. 157–165.

48. Hall, A.M. and E.D. Roberson, Mouse models of Alzheimer’s disease. Brain research bulletin, 2012. 88(1): p. 3–12.

49. Radde, R., et al., Aβ42-driven cerebral amyloidosis in transgenic mice reveals early and robust pathology. EMBO reports, 2006. 7(9): p. 940.

50. Maxwell, S.E., H.D. Delaney, and K. Kelley, Designing experiments and analyzing data: A model comparison perspective. 2017: Routledge.

51. Tukey, J.W., Comparing individual means in the analysis of variance. Biometrics, 1949: p. 99–114.

52. Lakens, D., Calculating and reporting effect sizes to facilitate cumulative science: a practical primer for t-tests and ANOVAs. Frontiers in psychology, 2013. 4: p. 62627.

53. Hosmer Jr, D.W., S. Lemeshow, and R.X. Sturdivant, Applied logistic regression. 2013: John Wiley & Sons.

54. Stone, M., Cross-validatory choice and assessment of statistical predictions. Journal of the royal statistical society: Series B (Methodological), 1974. 36(2): p. 111–133.

55. Hanley, J.A. and B.J. McNeil, The meaning and use of the area under a receiver operating characteristic (ROC) curve. Radiology, 1982. 143(1): p. 29–36.

